# SETD2 deficiency impairs β-catenin destruction complex to facilitate renal cell carcinoma formation

**DOI:** 10.1101/2020.07.13.200220

**Authors:** Hanyu Rao, Xiaoxue Li, Min Liu, Jing Liu, Wenxin Feng, Jin Xu, Wei-Qiang Gao, Li Li

**Affiliations:** State Key Laboratory of Oncogenes and Related Genes, Renji-Med X Clinical Stem Cell Research Center, Ren Ji Hospital, School of Medicine and School of Biomedical Engineering, Shanghai Jiao Tong University, Shanghai, 200127, China; School of Biomedical Engineering and Med-X Research Institute, Shanghai Jiao Tong University, Shanghai, China

## Abstract

Clear cell renal cell carcinoma (ccRCC) is a largely incurable disease that is highly relevant to epigenetic regulation including histone modification and DNA methylation. SET domain–containing 2 (SETD2) is a predominant histone methyltransferase catalyzing the trimethylation of histone H3 Lysine 36 (H3K36me3) and its mutations are highly relevant to clear cell renal cell carcinoma (ccRCC). However, its physiology role in ccRCC remains largely unexplored. Here we report that Setd2 deletion impairs the β-catenin destruction complex to facilitate ccRCC formation in a c-MYC-generated polycystic kidney disease (PKD) model, which can be relieved by an inhibitor of β-catenin-responsive transcription. Clinically, SETD2 loss is widely observed in ccRCC samples, and negatively correlated with expression of some members of β-catenin destruction complex, but positively correlated with the activation of Wnt/β-catenin signaling. Our findings thus highlight a previously unrecognized role of SETD2-mediated H3K36me3 modification in regulation of Wnt/β-catenin pathway in ccRCC.

**Summary:** Our findings for the first time reveal a previously unrecognized role of the SETD2-mediated H3K36me3 modification in regulation of the Wnt/β-catenin pathway in ccRCC and shed light on the molecular mechanisms underlying the formation of renal cell carcinoma with epigenetic disorders.

## Introduction

Renal cell carcinoma (RCC), originated from the renal tubular epithelium, is the second leading cause of death among all types of urologic cancer (Hsieh et al., 2017, Siegel et al., 2017, Rini et al., 2009). The most common subtype of RCC is clear cell renal cell carcinoma (ccRCC), which accounts for 75–80% of all diagnosed cases (Moch et al., 2016). Epigenetic regulation plays an important role in the initiation, development and treatment of malignant tumors including RCC (Dalgliesh et al., 2010, Baylin and Jones, 2011). Alterations in histone methylation are reported in several types of cancers and have also been examined in ccRCC (Shenoy et al., 2015). Recent studies have revealed that histone methylation plays a crucial role in epigenetic regulation and its mutation or functional loss as well as subsequent dysregulated downstream signaling facilitates oncogenic processes (Chen et al., 2020).

SETD2 is the only histone H3 lysine 36 histone (H3K36) methyltransferase that can alter the trimethylation status of H3K36 (H3K36me3) and regulates protein structures as well as its function (Mikkelsen et al., 2007, Hu et al., 2010). SETD2-induced H3K36me3 is a multifunctional histone marker associated with actively transcribed regions and is critical for many physiological processes, such as transcriptional regulation, chromosome segregation, DNA damage repair and alternative splicing (Li et al., 2013, Pfister et al., 2015, Zhang et al., 2014, Kanu et al., 2015). SETD2 was first linked to ccRCC when initial studies showed inactivation of the histone methyltransferase gene SETD2 as a common event in ccRCC cells (Varela et al., 2011, Pena-Llopis et al., 2012, Duns et al., 2010). Clinically, mutations of the SETD2 gene occur in up to 12% of early-stage ccRCC and result in loss of H3K36me3 in ccRCC-derived cells and tumors (Simon et al., 2014, Hacker et al., 2016). The expression level of SETD2 is associated with the tumor size, clinical stage and risk of carcinoma-related death, suggesting the worse prognosis in ccRCC patients with SETD2 loss of function (Liu et al., 2015, Ho et al., 2016a). As a tumor suppressor, SETD2 has been increasingly identified as a common mutation across cancer types, including lung cancer (Walter et al., 2017, Lee et al., 2019), intestinal cancer (Yuan et al., 2017), glioma (Fontebasso et al., 2013), gastrointestinal tumors (Yuan et al., 2017, Huang et al., 2016), osteosarcoma (Sakthikumar et al., 2018) and leukemia (Zhu et al., 2014, Skucha et al., 2018). Our previous studies have also reported that SETD2 is pivotal for bone marrow mesenchymal stem cells differentiation (Wang et al., 2018), maternal epigenome, genomic imprinting and embryonic development (Xu et al., 2019), sperm development (Zuo et al., 2018), and V(D)J recombination in normal lymphocyte development (Ji et al., 2019). SETD2 loss leads to pancreatic carcinogenesis (Niu et al., 2019). Mutations or functional loss of SETD2 produces dysfunctional tumor tissue proteins, causing tumorigenesis, progression, chemotherapy resistance and unfavorable prognosis (Chen et al., 2020). However, the mechanistic characterization of SETD2 in renal tumorigenesis and the involvement of particular signaling pathways remain elusive.

Numerous tumor suppressor genes have been reported to be partially or completely silenced due to hypermethylation. In particular, some members of the Wnt/β-catenin signaling pathway have been shown to be epigenetically silenced in several studies in ccRCC, such as SFRPs (Shenoy et al., 2015, Kawakami et al., 2011, Hirata et al., 2011). The activity of Wnt/β-catenin signaling within the process of tumorigenesis is tightly controlled by the phosphorylation, ubiquitination and degradation of β-catenin (Nick Barker, 2006), which is regulated by a highly processive “β-catenin destruction complex” in the pathogenesis of renal cancer including Adenomatosis Polyposis Coli (APC), Glycogen Synthase Kinase 3 Beta (GSK3β) and Beta-Transducin Repeat Containing E3 Ubiquitin Protein Ligase (BTRC) (Banumathy and Cairns, 2010). However, it remains unclear whether epigenetic regulators act epigenetically to regulate Wnt/β-catenin signaling through the β-catenin destruction complex in the formation of ccRCC.

Here, we established mouse models to determine the importance of Setd2 during the development of renal tubules and formation of ccRCC in a Setd2 deletion model alone or in a combination with a c-MYC-generated PKD model. Our RNA-seq and ChIP-seq analysis revealed that the Setd2 deficiency resulted in hyperactivation of Wnt/β-catenin signaling through down-regulating its negative regulators including APC, GSK3β and BTRC. Such negative correlation was also observed in clinical ccRCC samples. In addition, inhibition of the β-catenin-responsive transcription relieved the symptom caused by Setd2 deletion in mice. Our findings, for the first time, demonstrated a previously unrecognized role of the Setd2-mediated H3K36me3 modification in regulation of the Wnt/β-catenin pathway in ccRCC. Such novel autochthonous mouse model of ccRCC driven by c-MYC and deficiency of Setd2, will be useful for pre-clinical researches on ccRCC with epigenetic disorders.

## Results

### Setd2 deletion induces aberrant development of renal tubules in mice

To explore a possible role of SETD2 in kidney development and tumorigenesis in vivo, we generated Setd2-floxed mice and deleted the Setd2 gene in the kidney using a transgenic Ksp1.3/Cre mouse line (Shao et al., 2002) which has been widely utilized to model kidney cancer in mice (Harlander et al., 2017, Nargund et al., 2017). The expression of Cre from the Ksp-Cre is driven by the kidney-specific Cadherin 16 promoter, which begins expression at embryonic day 14.5 in epithelial cells of the developing kidney and continues to be expressed in tubular epithelial cells in adults (Shao et al., 2002). The resulting Ksp^Cre+^; Setd2^flox/flox^ mice are hereafter referred as Setd2^-KO^ mice. Though inactivation of Setd2 profoundly reduced H3K36me3 levels within tubular epithelial cells (Figure 1A and B), over a 5-month period, Setd2^-KO^ mice were viable and fertile and exhibited no gross phenotypic abnormalities compared to their control littermates (Figure 2A-C). However, after STZ treatment (150mg/kg), an agent that is toxic to kidneys in mammals and well-used to induce RCC (Hard, 1985, Yang et al., 2016), Setd2^-KO^ mice exhibited distinct aberrant development of renal tubules over a period of 4-week observations compared to their control littermates (Figure 1A). Moreover, Setd2^-KO^ mice treated with STZ showed enlarged aberrant areas in their kidneys (Figure 1B), and increased levels of blood urea nitrogen (BUN) and creatinine compared to STZ treated WT mice (Figure 1C), indicating that loss of Setd2 resulted in renal tubular dysfunction. When the survival of 20 mice (10 Setd2^fl/fl^, 10 Setd2^-KO^) after STZ treatment was monitored, a markedly increased mortality in Setd2^-KO^ mice compared with their control littermates was observed (Figure 1D). Interestingly, over a 10-month period, though comparable volumes of kidneys were observed in Setd2^-KO^ and Setd2^fl/fl^ mice, Setd2^-KO^ mice exhibited excessive growth of epithelial cells and aberrant development of renal tubules (Figure 2A-C). These results demonstrate that Setd2 deficiency results in dysfunction and aberrant development of renal tubules, but it is not sufficient to induce kidney tumors.

**Figure 1.**
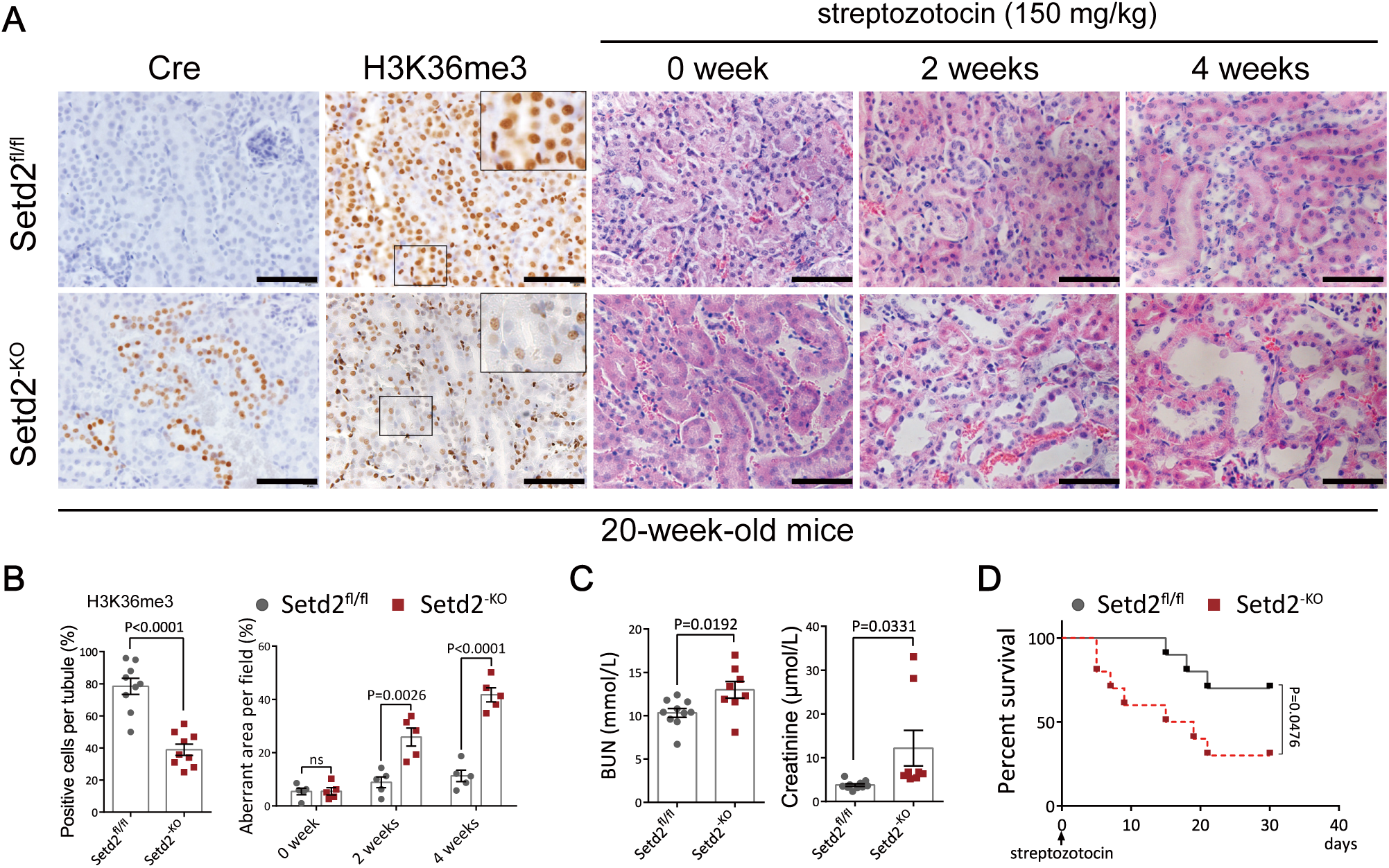
Setd2 deletion induces aberrant development of renal tubules. **(A)** Representative IHC images showing protein expression upon Setd2 deficiency. And weekly serial H&E of STZ treated kidney sections from Setd2^fl/fl^ and Setd2^-KO^ mice. **(B)** Metrics for IHC and H&E assays in kidney sections from STZ treated Setd2^fl/fl^ and Setd2^-KO^ mice. **(C)** Metrics for blood Urea Nitrogen and creatinine levels from STZ treated Setd2^fl/fl^ and Setd2^-KO^ mice. **(D)** Kaplan-Meier survival curve of Setd2^fl/fl^ and Setd2^-KO^ mice treated with STZ. Scale bars: 80 μm in A. Statistical comparisons were made using a 2-tailed Student’s t test. Data are represented as mean ± SEM.

**Figure 2.**
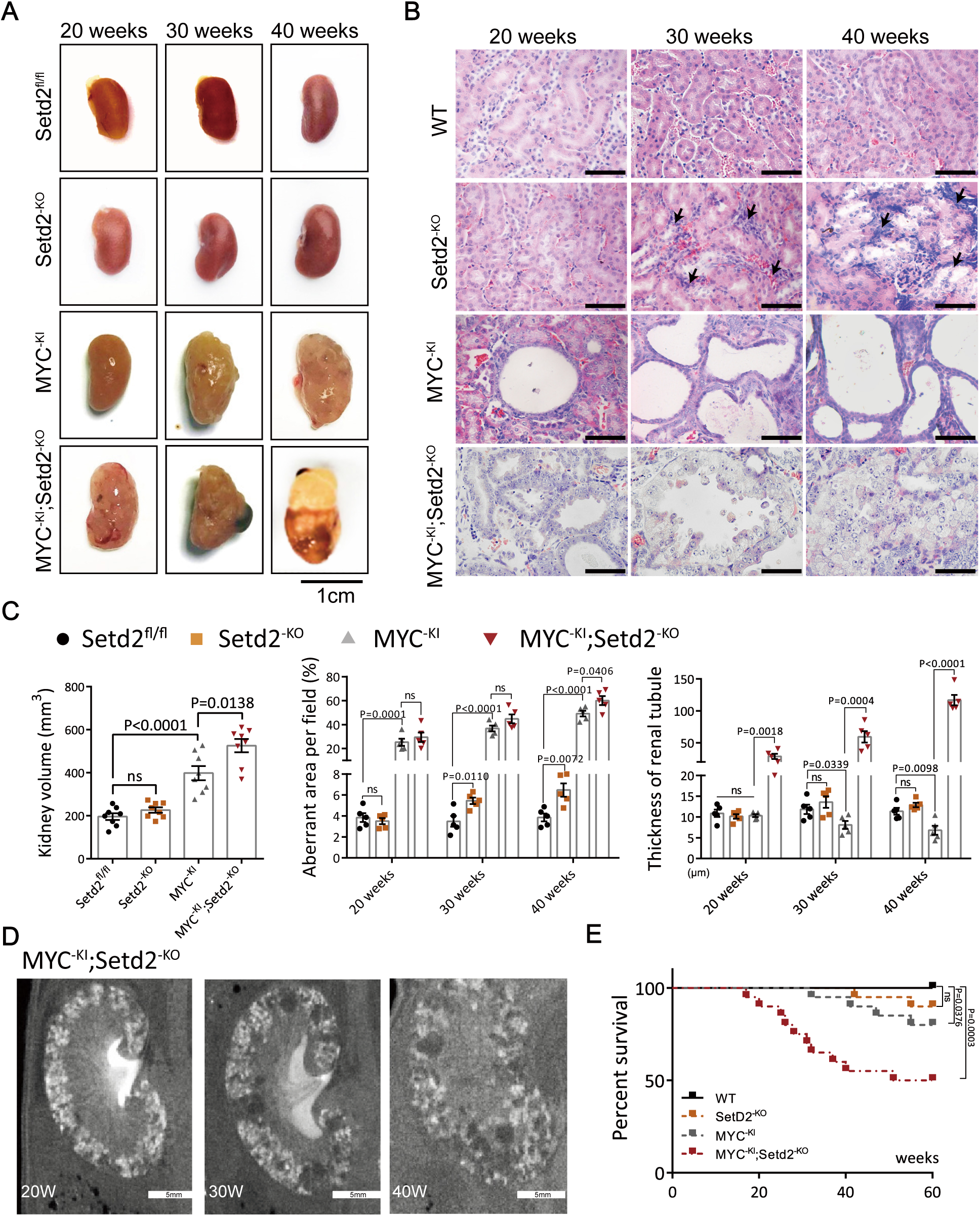
Setd2 deletion accelerates tumor formation in c-MYC-transgene mice. **(A)** Kidney volumes of Setd2^fl/fl^, Setd2^-KO^, MYC^-KI^ and MYC^-KI^; Setd2^-KO^ mice. **(B)** Weekly serial H&E of kidney sections following c-MYC activation and Setd2 deficiency. **(C)** Metrics for Kidney volumes and H&E assays in kidney sections from Setd2^fl/fl^, Setd2^-KO^, MYC^-KI^ and MYC^-KI^; Setd2^-KO^ mice. **(D)** Example of longitudinal μCT imaging of tumor development in a MYC^-KI^; Setd2^-KO^ mouse. **(E)** Kaplan-Meier survival curve of Setd2^fl/fl^, Setd2^-KO^, MYC^-KI^ and MYC^-KI^; Setd2^-KO^ mice. Scale bars: 1cm in A; 80 μm in B; 5mm in D. Statistical comparisons were made using a 2-tailed Student’s t test. Data are represented as mean ± SEM.

### Setd2 inactivation promotes ccRCC formation when combined with a c-MYC-generated PKD model

It has been demonstrated that elevated c-MYC activity can induce renal cancer using transgenic mice expressing a stable exogenous c-MYC in renal epithelial cells (Shroff et al., 2015, Bailey et al., 2017). Thus, to explore whether Setd2 loss can enhance c-MYC mediated RCC, we established our own c-MYC-transgene mouse line by overexpressing the c-MYC under the control of the Ksp promoter (Ksp^Cre+^; MYC^R26StopFL/+^ mice, hereafter referred as MYC^-KI^). Ectopic expression of c-MYC mediated by Ksp-Cre was observed within renal tubular epithelial cells (Supplemental Figure 1A). Surprisingly, these mice only exhibited enlarged kidney size and renal cysts, which are histologically associated with polycystic kidney disease (PKD) (Figure 2A-C). No apparent tumors were detected in MYC^-KI^ mice until 10 months. This data suggested that ectopic expression of c-MYC driven by Ksp1.3/Cre does not generate distinct tumors, which may due to its particular driven promoter or induction model. To determine if SETD2 deletion would play a synergistic effect on induction of ccRCC in c-MYC-generated PKD model, we generated Ksp^Cre+^; MYC^R26StopFL/+^ Setd2^flox/flox^ mice (hereafter referred as MYC^-KI^; Setd2^-KO^ mice). Ksp1.3/Cre mediated deletion of Setd2 profoundly reduced H3K36me3 levels within renal tubular epithelial cells (Supplemental Figure 1A). Importantly, after a 10-month period, larger volume of kidneys was observed containing distinct neoplastic masses in MYC^-KI^; Setd2^-KO^ mice compared to MYC^-KI^ mice (Figure 2A-C). MYC^-KI^; Setd2^-KO^ mice exhibited obvious structural abnormalities characterized by an unceasing, abnormal and excessive proliferation of renal tubular epithelial cells which gradually overfilled the tubule, whereas these phenotypes were not observed in MYC^-KI^ mice (Figure 2B-C). To follow up these observations in mice, we then monitored the formation of tumors in MYC^-KI^; Setd2^-KO^ mice using contrast-assisted micro computed tomography (μCT) imaging (Figure 2D). We found that tumor numbers and sizes in MYC^-KI^; Setd2^-KO^ mice increased proportionally within 20–40 weeks in a time-dependent manner. Moreover, as shown by Kaplan-Meier survival plots, their life span was much shorter compared to MYC^-KI^ mice (Figure 2E). These results demonstrate that Setd2 deletion accelerates the onset and increases the incidence of tumor formation.

Immunohistochemical staining revealed that the tumor cells in MYC^-KI^; Setd2^-KO^ mice displayed central features of human ccRCC, including clear cytoplasm and positive membranous staining of carbonic anhydrase IX (CA9), a major marker to diagnose ccRCC (Figure 3A) (Fu et al., 2011). Setd2^-KO^ and MYC^-KI^; Setd2^-KO^ mice exhibited excessive proliferation of renal tubular epithelial cells compared to Setd2^fl/fl^ and MYC^-KI^ mice as measured by Ki-67 staining. Stronger lipid accumulation, which is frequently observed in ccRCC (Han et al., 2016), were detected in the “clear” cells of MYC^-KI^; Setd2^-KO^ mice compared to MYC^-KI^ mice (Figure 3A and B). To compare these mouse tumors with human ccRCC, we performed gene expression profile analysis of these mouse tumors in comparison to human TCGA clear cell RCC (KIRC) kidney cancers. Correlation analysis of the average expression values for all unique orthologous gene pairs between human ccRCC and mouse ccRCC revealed a strong correlation in global transcriptional profiles (Figure 3C). Next, we performed stainings for lotus tetragonolobus lectin (LTL) that marks proximal convoluted tubule and for Tamm-Horsfall protein (THP) that marks distal convoluted tubule. We found that, MYC^-KI^; Setd2^-KO^ tumors were positive for THP but not LTL (Figure 3D), suggesting that these tumors originated from the distal tubule. Together, these results demonstrate that SETD2 deficiency accelerates ccRCC development in MYC^-KI^ mice and support the notion that SETD2 is a tumor suppressor in kidney.

**Figure 3.**
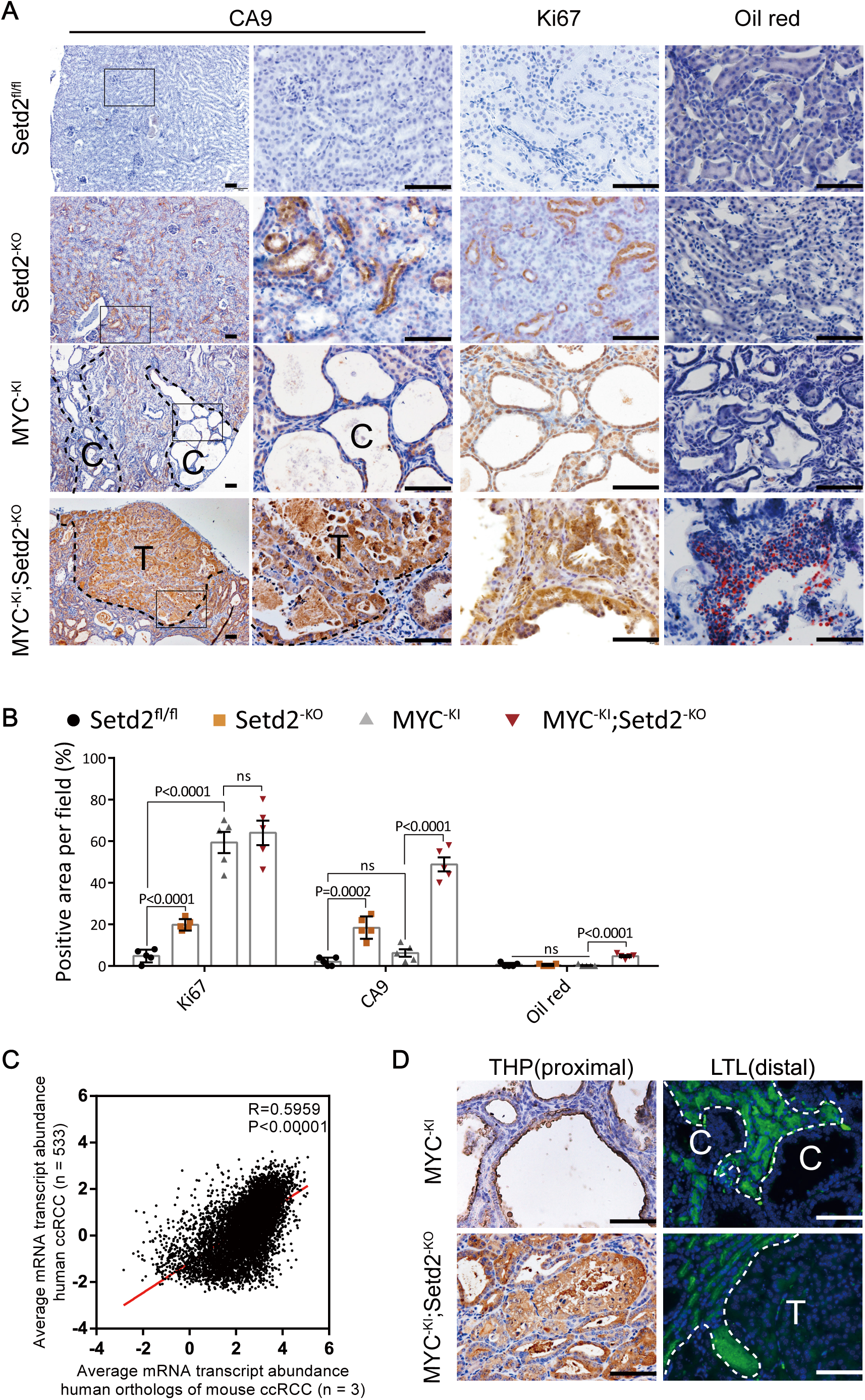
Setd2 deletion and c-MYC overexpression allows the evolution of ccRCC in mice. **(A)** Representative IHC images of CA9, Ki67 and Oil red of Setd2^fl/fl^, Setd2^-KO^, MYC^-KI^ and MYC^-KI^; Setd2^-KO^ kidneys. **(B)** Metrics for IHC assays in kidney sections from Setd2^fl/fl^, Setd2^-KO^, MYC^-KI^ and MYC^-KI^; Setd2^-KO^ mice. **(C)** Sample-averaged, normalized and log-transformed gene expression values for all unique orthologous gene pairs in human and mouse ccRCC (black dots). (Pearson correlation test, P < 1 × 10-16). **(D)** Representative IHC images of THP and LTA of MYC^-KI^ and MYC^-KI^; Setd2^-KO^ kidneys. Scale bars: 80 μm in A and D. Statistical comparisons were made using a 2-tailed Student’s t test. Data are represented as mean ± SEM.

To provide additional supporting evidence for the synergistic effect on tumor formation by SETD2 deletion, we then performed experiments to delete or over-express Setd2 using lentivirus-mediated knockout or overexpression systems in WT and c-MYC-overexpressed renal primary tubular epithelial cells (PTECs), which are generally regarded as the normal counterparts of ccRCC (Li et al., 2016). In this way, we were able to examine more detailed biological features related to tumorigenesis in PTECs. As shown in Supplemental Figure 2A and 3A, knockout of Setd2 led to a significant decrease at Setd2 mRNA level while overexpression of Setd2 and c-MYC caused a significant increase at their mRNA levels. Notably, PTECs^Setd2-KO^ and PTECs^MYC-KI;Setd2-KO^ resulted in more intensive proliferation, migration and invasion capacity compared to PTECs^Vecotr^ and PTECs^MYC-KI^ (Supplemental Figure 2B and 2C). On the other hand, overexpression of SETD2 in PTECs^Vecotr^ and PTECs^MYC-KI^ resulted in attenuated proliferation, migration and invasion capacity compared to their negative controls (Supplemental Figure 3B and 3C).

### Setd2 deficient renal tubular epithelial cells display hyperactive Wnt/β-catenin Signaling

To gain an insight into the mechanism of how Setd2 ablation promotes ccRCC, we performed RNA-seq using PTECs isolated from 20-week-old Setd2^fl/lf^ and Setd2^-KO^ mice. We found that the global transcriptome was changed dramatically in PTECs^Setd2-KO^ (Figure 4A). Gene ontology (GO) analysis indicated that Setd2 loss significantly enriched the genes associated with cellular metabolism, cell cycle and mitosis, cell proliferation, migration and cell-cell adhesion (Supplemental Figure 4A). To better understand Setd2-mediated signal circuits, we performed gene set enrichment analysis (GSEA) and found that Setd2 loss significantly enriched the genes linked to cell metabolism and tight junction (Figure 4B). Then, we performed RNA-seq using the PTECs isolated from 20-week-old MYC^-KI^ and MYC^-KI^; Setd2^-KO^ mice (Figure 4C). GSEA analysis revealed that Setd2-regulated genes coincided with the Wnt/β-catenin signaling signatures (Figure 4D). RT-qPCR analysis verified that the expression levels of Wnt-induced target genes, such as c-Myc, Axin2 and Ccnd1, were up-regulated in PTECs^Setd2-KO^ and PTECs^MYC-KI;Setd2-KO^ compared to control cells (Figure 4E). Moreover, accumulations of activated β-catenin and Ccnd1 were increased in MYC^-KI^; Setd2^-KO^ mice compared to MYC^-KI^ mice (Figure 4F and G). These data indicated that Wnt/β-catenin signaling was activated after the deletion of Setd2. In addition, other signaling pathways were also activated, including the Notch signaling pathway, VEGF signaling pathway, MAPK signaling pathway and TGF-β signaling pathway based on GSEA analysis (Supplemental Figure 5A). However, there were not significant differences for expression levels of mTOR, PTEN and SMADs in the two groups (Supplemental Figure 5B). Together, these results indicate that Setd2 deficiency leads to Wnt/β-catenin activation and metabolic disturbance, which promotes the cell proliferation and tumorigenesis of renal tubular cells in ccRCC.

**Figure 4.**
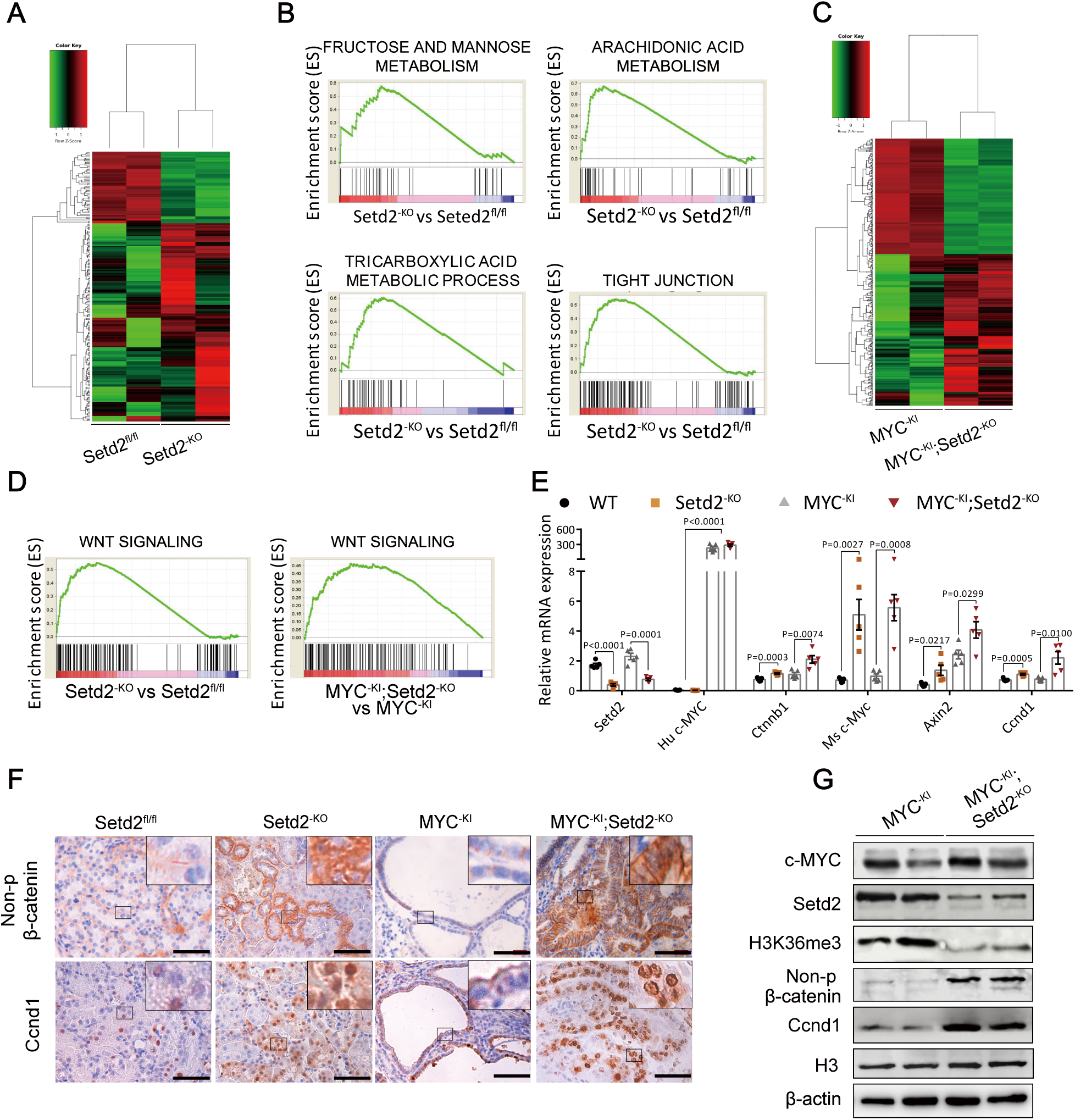
Setd2 deficient renal tubular epithelial cells display hyperactive Wnt/β-catenin signaling. **(A)** Heatmap of genes with significantly different expression in age-matched Setd2^fl/fl^ and Setd2^-KO^ renal tubular epithelial cells based on unsupervised hierarchical agglomerative clustering. **(B)** GSEA enrichment plots of differentially expressed genes associated with Setd2 deletion. **(C)** Heatmap of genes with significantly different expression in age-matched MYC^-KI^ and MYC^-KI^; Setd2^-KO^ renal tubular epithelial cells based on unsupervised hierarchical agglomerative clustering. **(D)** GSEA enrichment plots of differentially expressed genes belonging to Wnt/β-catenin signaling pathway associated with Setd2 deletion. **(E)** RT-qPCR analysis of β-catenin target genes in Setd2^fl/fl^, Setd2^-KO^, MYC^-KI^ and MYC^-KI^; Setd2^-KO^ kidneys. **(F)** IHC images of active β-catenin and Ccnd1 in Setd2^fl/fl^, Setd2^-KO^, MYC^-KI^ and MYC^-KI^; Setd2^-KO^ tumors. **(G)** Immunoblotting analysis in MYC^-KI^ cysts and MYC^-KI^; Setd2^-KO^ tumors with antibodies indicated on the left. Scale bars: 80 μm in G. Statistical comparisons were made using a 2-tailed Student’s t test. Data are represented as mean ± SEM.

### Setd2-mediated H3K36me3 is essential for transcription of members of “β-catenin destruction complex”

To further understand the underlying mechanisms and identify genes directly regulated by Setd2 and H3K36me3 at a genome-wide scale, we performed chromatin immunoprecipitation experiments followed by next-generation sequencing (ChIP-seq) assays using a H3K36me3-specific antibody in PTECs isolated from Setd2^-KO^ and Setd2^fl/fl^ mice. In line with our hypothesis, H3K36me3 peaks were enriched around transcription area, and Setd2 deletion resulted in the reduction of H3K36me3 peaks in PTECs (Figure 5A and B). Among a total of 19,032 genes, 780 genes were differentially expressed in PTECs^Setd2-KO^. A total of 2 723 and 12 199 H3K36me3 peaks were identified in PTECs^Setd2-KO^ and control cells respectively. We next analyzed H3K36me3 peaks that were absent in PTECs^Setd2-KO^, and found those peaks were from 3 420 genes (Figure 5C). Among them, there were several important inhibitors of the Wnt/β-catenin signaling pathway including APC, PTPA, CSNK1A1, CSNK1D, GSK3α, GSK3β, BTRC, COPS5, JADE1, JADE2, CEL1 and CUL2. To demonstrate the involvement of Setd2 and H3K36me3 histone markers in these genes, we performed ChIP-qPCR to validate H3K36me3 occupancies in the transcription area of these members of “β-catenin destruction complex”. The H3K36me3 coding for these genes were presented in figures 5D, which indicated that Setd2 loss eliminated H3K36me3 modifications in their transcription area. To further investigate the transcription status in these genes, we performed ChIP-qPCR analysis using antibodies specific for RNA polymerase II (Pol II). Focusing on the potential candidates implicated in β-catenin destruction, we noticed that APC, GSK3B and BTRC were listed as the top candidate, displaying significantly less H3K36me3 modifications and enrichment of Pol-II at their gene body-retained locus upon Setd2 deletion (Figures 5E). RT-qPCR analysis of pre-selected genes showed that expression levels of both pre-mRNA and mRNA of APC, GSK3β and BTRC were downregulated in Setd2 deficient mice (Figure 5F). Together, our results demonstrate that Setd2-mediated H3K36me3 in the transcription area of some members of “β-catenin destruction complex” is important for their transcription and Wnt/β-catenin signaling in mice.

**Figure 5.**
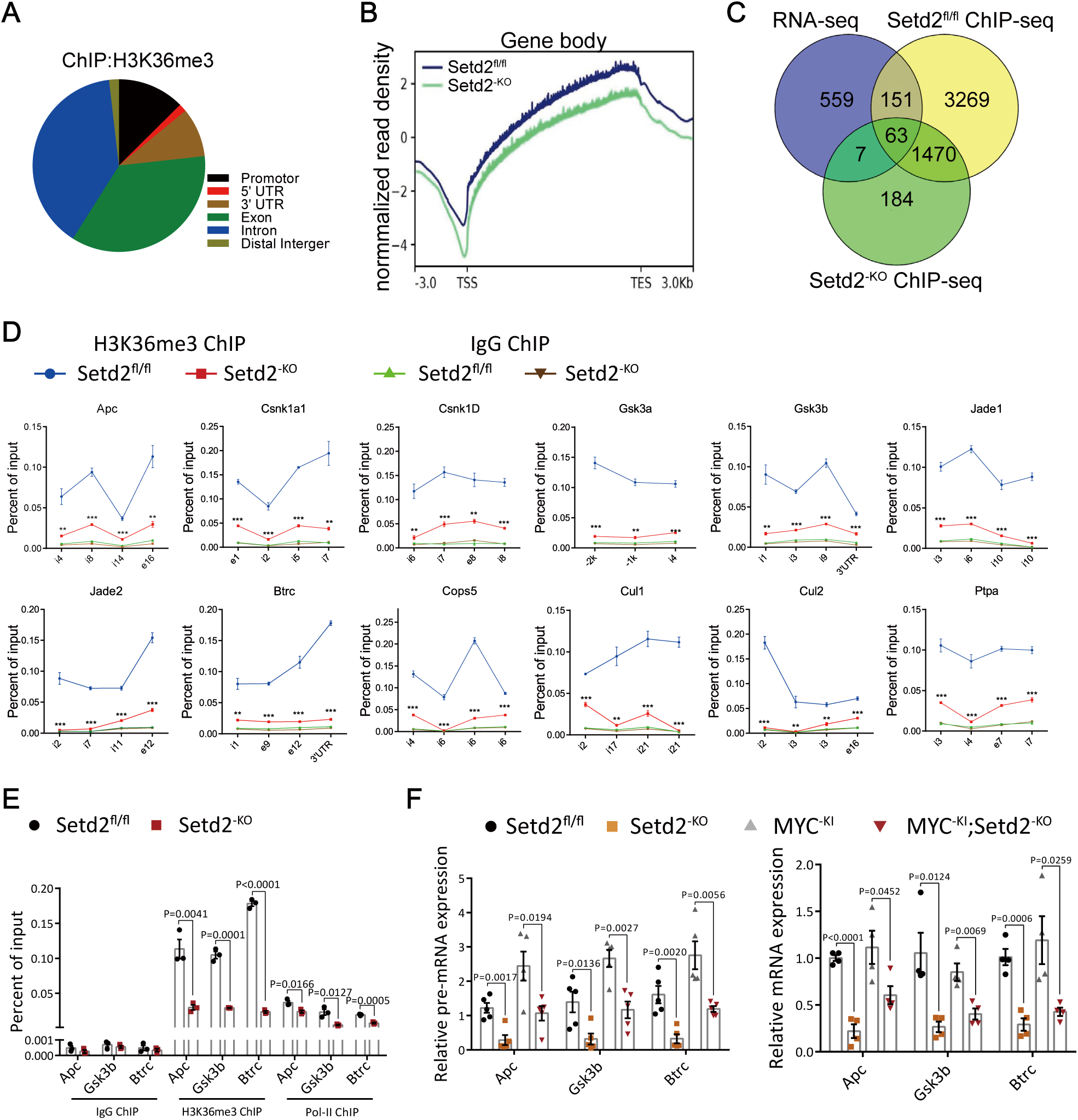
Setd2-mediated H3K36me3 is essential for the transcription of members of “β-catenin destruction complex”. **(A)** Analysis of the occupancy of H3K36me3 ChIP-seq peaks in gene bodies and intergenic regions. **(B)** Normalized read density of H3K36me3 ChIP-seq signals of Setd2^fl/fl^ and Setd2^-KO^ kidneys from 3 kb upstream of the TSS to 3 kb downstream of the TES. **(C)** Venn diagram illustration of H3K36me3 peaks of Setd2^fl/fl^ and Setd2^-KO^ kidneys, and their overlap with differential expression genes determined by RNA sequencing. **(D)** ChIP-qPCR analysis of H3K36me3 occupancy to gene body-retained locus, using IgG as the control. **(E)** ChIP-qPCR analysis of H3K36me3 and Pol-II occupancy to gene body-retained locus, using IgG as the control. **(F)** RT-qPCR analysis of pre-mRNA and mRNA levels of Apc, Gsk3b and Btrc in Setd2^fl/fl^, Setd2^-KO^, MYC^-KI^ and MYC^-KI^; Setd2^-KO^ kidneys. Statistical comparisons were made using a 2-tailed Student’s t test. Data are represented as mean ± SEM.

### SETD2 activity is negatively correlated with Wnt/β-catenin signaling and ccRCC tumorigenesis

To determine the clinical functional relevance between SETD2 and Wnt/β-catenin signaling, we analyzed transcriptomic signature in ccRCC patients (TCGA-KIRC database, n=606). Clinically, mutations or low expression of SETD2 were widely observed in ccRCC samples (Figure 6A). Following an increased mutation rate of SETD2 gene, expression levels of APC, GSK3β and BTRC were decreased significantly with the progression of the tumor (Figure 6B). Moreover, there were also positive correlations between the mRNA levels of SETD2 and that of APC, GSK3β and BTRC respectively (Figure 6C). Thus, mutations or low expression of SETD2 were clinically related to the downregulations of APC, GSK3β and BTRC in ccRCC. CA9 and VIM are markers of ccRCC, while c-MYC and CCND1 are Wnt/β-catenin target genes. In accordance to our preceding results, analyses of transcriptomic signature showed inverse correlations between the mRNA level of SETD2 and that of CA9, VIM, c-MYC and CCND1 (Figure 6D). Therefore, activity of SETD2 is negatively correlated with the activation of Wnt/β-catenin signaling and ccRCC tumorigenesis in ccRCC samples. Collectively, these findings indicate that the deficiency of SETD2 is clinically related to the downregulations of the members of “β-catenin destruction complex”, which leads to the activation of Wnt/β-catenin signaling and ccRCC tumorigenesis.

**Figure.6.**
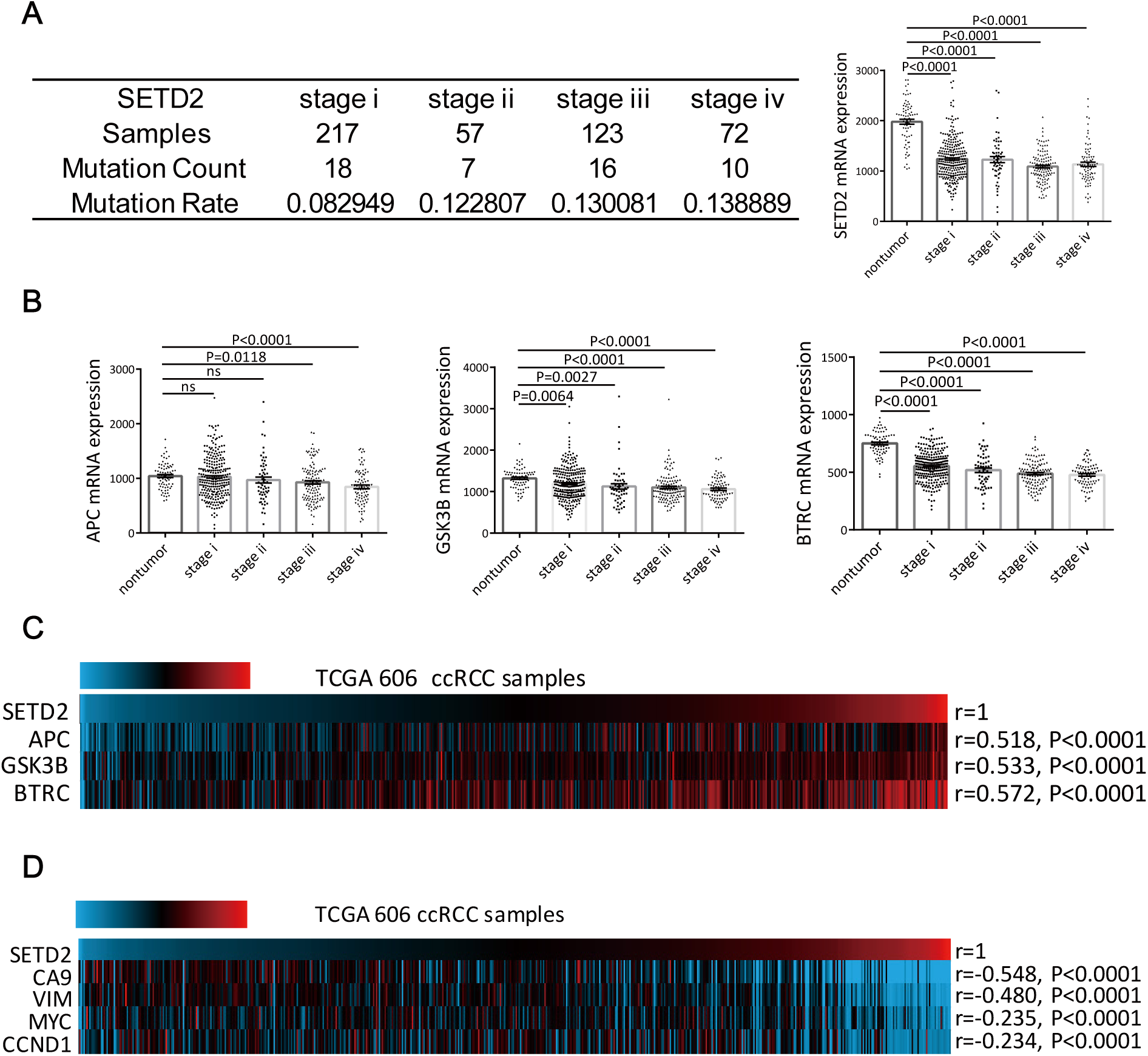
SETD2 is highly relevant to ccRCC and is essential for expression of APC, GSK3β and BTRC in ccRCC clinical samples. **(A)** Mutation rate and expression level of SETD2 in ccRCC stages. **(B)** Expression levels of APC, GSK3β and BTRC in ccRCC stages. **(C)** Correlations between the expression levels of SETD2 gene and that of APC, GSK3β and BTRC respectively in KIRC samples. (Pearson correlation test, n=606). **(D)** Correlations between the expression levels of SETD2 gene and ccRCC markers and Wnt/β-catenin target genes in KIRC samples. (Pearson correlation test, n=606). Statistical comparisons were made using a 2-tailed Student’s t test. Data are represented as mean ± SEM.

### Inhibition of Wnt/β-catenin pathway relieves the symptom caused by Setd2 deletion in mice

Given that Setd2 loss-mediated tumorigenesis is dependent on hyperactivation of Wnt/β-catenin signaling, we next investigated whether this model is sensitive to Wnt/β-catenin signaling inhibitors. iCRT-14, a potent inhibitor of β-catenin-responsive transcription (Gonsalves et al., 2011), was utilized to silence β-catenin signaling. For preclinical therapeutic studies, daily intra-peritoneal injection of iCRT-14 was performed for a month on 40-week-old tumor-bearing mice (MYC^-KI^; SETD2^-KO^). After iCRT-14 treatment, as expected, stainings of activated β-catenin and Ccnd1 were decreased compared to DMSO-treated mice (Figure 7A). Moreover, iCRT-14-treated MYC^-KI^; Setd2^-KO^ mice displayed decreased proliferation of renal tubular epithelial cells compared to DMSO-treated mice as measured by Ki-67 staining (Figure 7A). And these tumors showed attenuated staining of CA9 and less lipid accumulation (Figure 7A), indicating that the ccRCC-related symptoms were relieved after the utilization of iCRT-14. In PTECs, iCRT-14 treatment dramatically attenuated proliferation, migration and invasion capacities of Setd2 deficient PTECs. (Figure 7B and C). Moreover, Wnt-induced target genes were also downregulated in iCRT-14 treated Setd2 deficient PTECs (Figure 7D). Collectively, these results demonstrate that Setd2 loss aggravates ccRCC progression in a Wnt/β-catenin signaling-dependent manner.

**Figure.7.**
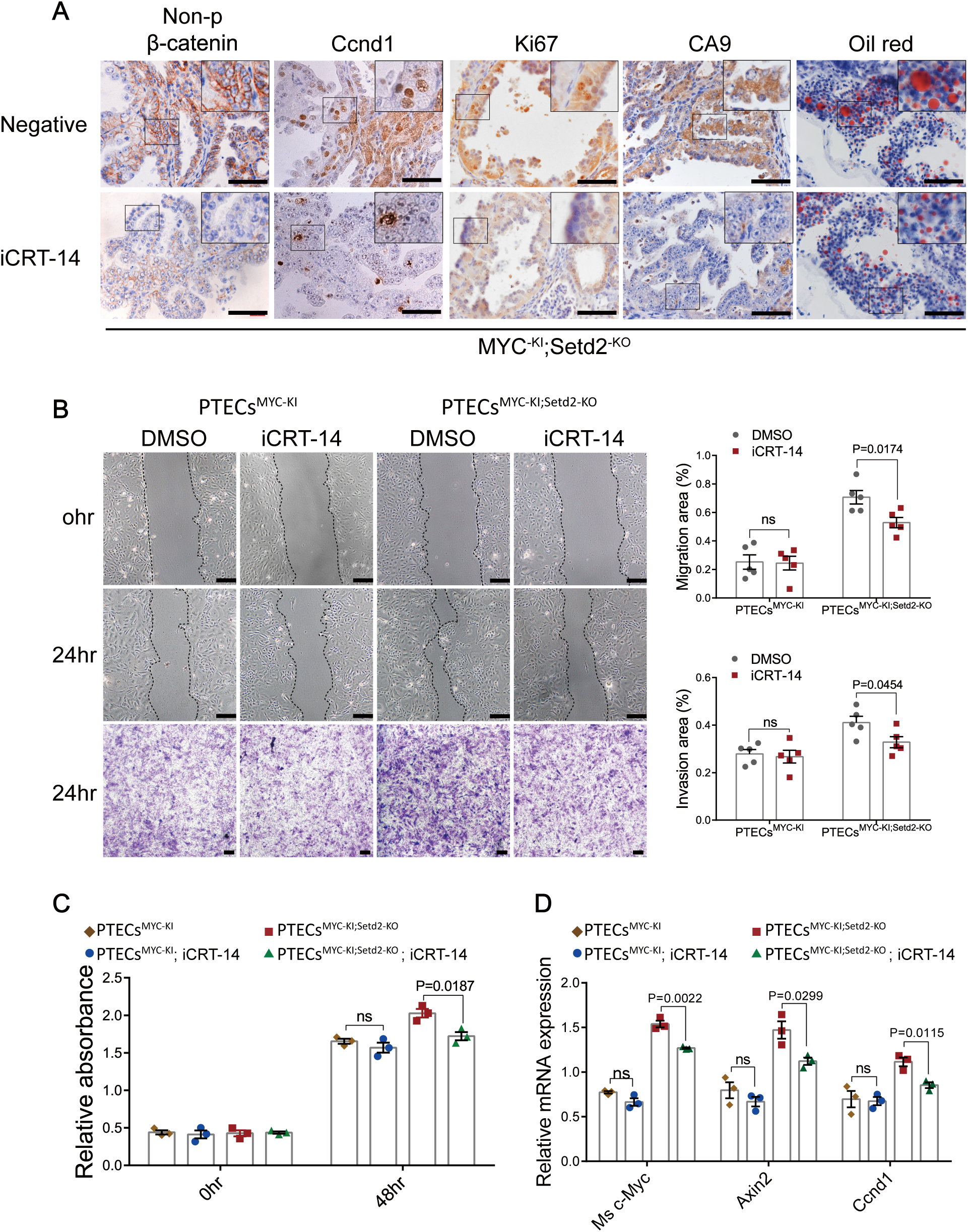
Inhibition of Wnt/β-catenin pathway relieves the symptom caused by Setd2 deletion in mice. **(A)** IHC images of active β-catenin, Ccnd1, Ki67, CA9 and Oil red in MYC^-KI^; Setd2^-KO^ tumors treated with iCRT-14 or negative control. **(B)** Cell migration and invasion abilities of PTECs were compared to their control cells, as demonstrated by cell wound scratch and trans-well assays. **(C)** Cell proliferation abilities of PTECs were compared to their control cells. **(D)** RT-qPCR analysis of mRNA levels in iCRT-14 treated PTECs. Scale bars: 80 μm in A and B. Statistical comparisons were made using a 2-tailed Student’s t test. Data are represented as mean ± SEM.

## Discussion

The present study demonstrates that mice with Setd2 knockout in renal epithelial cells, under the control of Ksp1.3/Cre, exhibit aberrant development of renal tubules, and that when combined with a c-MYC-driven PKD model, distinct ccRCC is generated. It has been previously shown that elevated c-MYC activity can induce renal cancer using transgenic mice expressing a stable ectopic c-MYC in renal epithelial cells (Shroff et al., 2015, Bailey et al., 2017). However, in current study, we show that under the control of Ksp1.3/Cre in vivo, ectopic expression of c-MYC can induce PKD only, which is probably due to its particular driven promoter or induction model. A combination of sustained ectopic expression of c-MYC with deficiency of SETD2 is required to generate ccRCC. Our work is consistent with recent clinical reports indicating that mutations of the SETD2 gene occur in up to 12% of early-stage ccRCC and the numbers of H3K36me3-positive nuclei are reduced an average of ∼20% in primary ccRCC (Simon et al., 2014, Hacker et al., 2016, Ho et al., 2016b), reinforcing the notion that SETD2-related epigenetic regulation plays a key role in ccRCC. Therefore, the c-MYC and Setd2 double mutant mice that we established may serve as a novel autochthonous genetically engineered mouse model of ccRCC for pre-clinical researches on ccRCC with epigenetic disorders. Toward this direction, it would be also applicable for basic and clinical researches on ccRCC to combine mutation of SETD2 gene with other frequently mutated tumor-suppressor genes, such as VHL, p53, PTEN, PBMR1 and RB1, which have been shown to cooperatively cause the evolution of RCCs and kidney diseases (Harlander et al., 2017, Albers et al., 2013, Nargund et al., 2017, Feng et al., 2015, Gossage et al., 2015).

Our work reveals a mechanism how SETD2 facilitates formation of ccRCC and a close clinical correlation between Setd2 mutation/downregulation and ccRCC development. At molecular level, we show that the Setd2 loss leads to down-regulation of the members of “β-catenin destruction complex” including Apc, Gsk3b and Btrc and consequently activates the Wnt/β-catenin signaling in both mice and PTECs. Notably, PTECs^Setd2-KO^ and PTECs^MYC-KI;Setd2-KO^ exhibit more intensive proliferation, migration and invasion capacity compared to PTECs^Vecotr^ and PTECs^MYC-KI^, respectively Clinically, we illustrate that SETD2 is frequently mutated or downregulated in ccRCC samples, and its activity is positively correlated with expression of APC, GSK3β and BTRC, but negatively correlated with the activation of Wnt/β-catenin signaling and ccRCC tumorigenesis. Such observations are further supported by our experiments showing that the symptom caused by Setd2 deletion can be relieved in the mutant mice. Therefore, our findings unveil for the first time an important role of the SETD2-mediated H3K36me3 modification in regulation of the Wnt/β-catenin pathway in ccRCC and provide insights into the molecular mechanisms underlying the formation of renal cell carcinoma with epigenetic disorders.

It is worth mentioning that SETD2 appears to interact with various proteins to influence transcription. Notably, SETD2-mediated H3K36me3 is reported to participate in cross-talks with other chromatin markers, including antagonizing H3K4me3 and H3K27me3 (Xu et al., 2019) and promoting de novo DNA methylation (Rondelet et al., 2016). In our study, we illustrate that expression of some members of “β-catenin destruction complex” is regulated by SETD2-mediated H3K36me3. However, little is known about the epigenetic functions of other chromatin markers on regulating these genes in ccRCC. It would be interesting to study the physiological role of the cross-talk between H3K36me3 and other chromatin markers in ccRCC in the future. In addition, recent studies have also revealed that SETD2 has a capacity to regulate cellular signaling and responses through modification of non-histone substrates, such as signal transducer and activator of transcription (STAT1) (Chen et al., 2017) and α-tubulin in some biological processes (Park et al., 2016), which are quite different from previous studies that have linked biological function of SETD2 to its methyltransferase activity on histones. In the present experiments, we showed that SETD2-mediated H3K36me3 promotes the expression of some members of “β-catenin destruction complex”. However, it remains to be determined whether SETD2 can interact with these proteins and methylate them in ccRCC.

Taken together, our findings have demonstrated the biological significance of SETD2 as a pivotal epigenetic regulator in ccRCC. The deficiency of SETD2 can promote the accumulation of β-catenin and result in hyperactivation of Wnt/β-catenin signaling. Clinically, SETD2 activity is positively correlated with the expression levels of some members of “β-catenin destruction complex”, and negatively correlated with ccRCC tumorigenesis. Furthermore, inhibition of the Wnt/β-catenin pathway relieves the symptom caused by Setd2 deletion in mice. We reveal for the first time a previously unacknowledged role of SETD2-mediated H3K36me3 modification in regulation of the Wnt/β-catenin pathway underlying ccRCC. We also establish an autochthonous mouse model of ccRCC driven by a combination of c-MYC overexpression and deficiency of SETD2, which will be useful for pre-clinical researches on ccRCC with epigenetic disorders. For clinical translation, pharmaceutical investigation of the cross-talks between SETD2-mediated H3K36me3 and WNT/β-catenin signaling may provide a potential promising strategy to ccRCC therapy.

## Materials and methods

### Mouse strains

SETD2^fl/fl^ mice were generated by Shanghai Biomodel Organism Co. using conventional homologous recombination in ES cells as previously reported (Niu et al., 2019). The Ksp^Cre^ mice (B6.Cg-Tg(Cdh16-cre)91Igr/J) and MYC^R26StopFL/+^ (C57BL_6N-Gt(ROSA)26Sor(tm13(CAG-MYC,-CD2)Rsky) were purchased from The Jackson Laboratory. SETD2^fl/fl^ mice were mated with Ksp^Cre^ mice to generate Ksp^Cre^; SETD2^flox/flox^ (SETD2^-KO^) mice in C57BL/6 background. MYC^R26StopFL/+^ mice were mated with Ksp^Cre^ mice to generate Ksp^Cre^; MYC^R26StopFL/+^ (MYC^-KI^) mice in C57BL/6 background. SETD2^-KO^ mice were mated with MYC^-KI^ mice to generate Ksp^Cre^; MYC^R26StopFL/+^ SETD2^flox/flox^ (MYC^-KI^; SETD2^-KO^) mice housing under same condition.

### *In vivo* mouse models

Experimental acute tubular injury model was induced by single-dose intravenous injection (i.v) of streptozotocin into indicated mice as previously described (Hard, 1985). Single-dose of streptozotocin (150 mg/kg body weight) was injected via the tail veil into 20-week-old SETD2^fl/fl^ and SETD2^-KO^ mice. Recipient mice were then harvested for kidney histological analysis 2-6 weeks after injection.

### BUN and creatinine test

Blood was collected from mice using tail vein blood collection method. Serum was analyzed for BUN and creatinine concentration by Shanghai Biomodel Organism Co. on a Beckman Coulter AU680 analyzer.

### Isolation of PTECs and cell cultures

The isolation procedures and phenotype identification were performed as previously described (Ark et al., 2013, Terryn et al., 2007). PTECs were maintained in Dulbecco’s modified Eagle’s medium/F-12 GLUTMAX-1 containing 10% fetal bovine serum (FBS), 100 U/ml of penicillin and 100 μg/ml of streptomycin (all used for cell culturing are from Sigma-Aldrich, St. Louis, MO). All the cells were maintained at 37°C under humidified air containing 5% CO2. Mycoplasma, bacteria, and fungi were tested as negative in these cultures.

### μCT imaging

Imaging of animals was performed as previously described (Almajdub et al., 2008). Contrast agents were manually injected intravenously through the tail catheter in 20–30s. A contrast agents was used: iodixanol (Visipaque, 320mg iodine/ml; GE Health Care, Oslo, Norway), a dimeric isomolar nonionic water-soluble radiographic contrast agent, molecular weight 1.6 kDa (iodine span 49.1%), osmolality 290 mOsm per kg water, used for the C57BL/6J studies; 2.5 gI/kg body weight was injected. A micro-CT system (Quantum GX2, PerkinElmer) with an X-ray source (focal spot size, 5 mm; energy range, 90 kV, 88 uA) were used. Mice were imaged every 10 weeks, beginning 20 weeks after birth.

### RNA isolation and quantitative RT-PCR

Total RNA was isolated from cultured cells or fresh samples with Trizol reagent (Invitrogen). cDNA was synthesized by reverse transcription using the Prime Script RT reagent kit (TaKaRa) and subjected to quantitative RT-PCR with SETD2, Mouse-c-Myc, Human-c-MYC, Ctnnb1, Axin2, Ccnd1, Apc, Gsk3b, Btrc and actin-beta or Gapdh primers in the presence of the SYBR Green Realtime PCR Master Mix (Thermo). Relative abundance of mRNA was calculated by normalization to actin-beta or Gapdh mRNA. Data were analyzed from three independent experiments and were shown as the mean ± SEM.

### Western blot analysis and antibodies

Cells were lysed with 100–300μL RIPA buffer supplemented with protease and phosphatase inhibitors (Millipore). The protein concentration was measured with the BCA Protein Assay (Bio-Rad). From each sample, 20–50 μg of total protein was separated by 8–12% SDS-PAGE gels and transferred onto nitrocellulose membranes (GM). Membranes were blocked in 5% BSA in TBS for 1 hour at room temperature, and then incubated with primary antibodies overnight at 4**°**C, washed in TBS containing 1% Tween20, incubated with an HRP-conjugated secondary antibody for 1 hour at room temperature, and developed by ECL reagent (Thermo). The immunoblots were quantified by Bio-Rad Quantity One version 4.1 software. Primary antibodies against SETD2 (LS-C332416), histone H3 (tri methyl K36) (ab9050) and CCND1 (Lot.51514) were purchased from Lifespan, Abcam, and Arigo biolaboratories. Antibodies against c-Myc (#13987), histone H3 (#9715), non-phospho (Active) β-Catenin (#8814) and actin-beta (#3700) were purchased from Cell Signaling Technology Inc.

### Histology and immunohistochemistry (IHC) Staining

Tissues were fixed in 10% buffered formalin and fixed tissues were sectioned for hematoxylin and eosin (H&E) staining. For IHC staining, paraffin-embedded tissues were deparaffinized, rehydrated, and subjected to a heat-induced epitope retrieval step by treatment with 0.01M sodium citrate (pH 6.0). Endogenous peroxidase activity was blocked with 0.3% (v/v) hydrogen peroxide in distilled water. The sections were then incubated with 0.3% Triton X-100 in PBS (137 mM NaCl, 2.7 mM KCl, 10 mM Na2HPO4, 2 mM KH2PO4, pH 7.4) for 15 minutes, followed by 10% goat serum in PBS for 1 hour. Subsequently, samples were incubated with Primary antibodies, diluted at appropriate proportion in 1% goat serum for 1 hour at 37°C. After three washes in PBS, sections were incubated with an HRP-conjugated secondary antibody for 1 hour at room temperature. Sections were counterstained with hematoxylin. Three random immunostaining images of each specimen were captured using a Leica DM2500 microscope and analyzed by Image-Pro Plus 6.0 software. Primary antibodies against CA9 (NB100-417), THP (#590C14A) and CCND1 (Lot.51514) were purchased from Novo Biologic, Arigo biolaboratories and ORiGene. Antibodies against Cre Recombinase (#15036) and non-phospho (Active) β-Catenin (#8814) actin-beta were purchased from Cell Signaling Technology Inc. Antibodies against Ki-67 (ab6526), histone H3 (tri methyl K36) (ab9050) and c-Myc (ab32072) were purchased from abcam. Lotus tetragonolobus lectin (LTL) was purchased from Alfa Aesar.

### RNA-seq and analyses

Kidney tissue from mice were harvested around 8-week for RNA preparation. NEB Next Ultra Directional RNA Library Prep Kit for Illumina (New England Biolabs, Ipswich, MA, USA) was used for the construction of sequencing libraries. The libraries were then subjected to Illumina sequencing with paired-ends 2×150 as the sequencing mode. The clean reads were mapped to the mouse genome (assembly GRCm38) using the HISAT2 software. Gene expression levels were estimated using FPKM (fragments per kilobase of exon per million fragments mapped) by StringTie. Gene annotation file was retrieved from Ensembl genome browser 90 databases. To annotate genes with gene ontology (GO) terms and KEGG pathways, ClusterProfiler (R package) was used. The functional enrichment analysis (GO and KEGG) were also performed with ClusterProfiler.

### ChIP-Seq and analyses

Cells were crosslinked with 1 % formaldehyde for 10 min at room temperature and quenched with 125 mM glycine. The fragmented chromatin fragments were pre-cleared and then immunoprecipitated with Protein A+G Magnetic beads coupled with anti-H2K36me3 (ab9050) antibody. After reverse crosslinking, ChIP and input DNA fragments were end-repaired and A-tailed using the NEBNext End Repair/dA-Tailing Module (E7442, NEB) followed by adaptor ligation with the NEBNext Ultra Ligation Module (E7445, NEB). The DNA libraries were amplified for 15 cycles and sequenced using Illumina NextSeq 500 with single-end 1×75 as the sequencing mode. Raw reads were filtered to obtain high-quality clean reads by removing sequencing adapters, short reads (length<50 bp) and low-quality reads using Cutadapt (v1.9.1) and Trimmomatic (v0.35). Then FastQC is used to ensure high reads quality. The clean reads were mapped to the mouse genome (assembly GRCm38) using the Bowtie2 (v2.2.6) software. Peak detection was performed using the MACS (v2.1.1) peak finding algorithm with 0.01 set as the p-value cutoff. Annotation of peak sites to gene features was performed using the ChIPseeker R package.

### Plasmids, transfection, and lentivirus

Setd2 cDNA and recombinase cDNA were generated by polymerase chain reaction and cloned into pCMV6-Entry vector with HA-tag. Then, the cDNA of Setd2 were cloned into the lentiviral expression vector pLenti.CMV.Puro.DEST. All the constructs generated were confirmed by DNA sequencing. For transfection, PTECs were transfected with the jetPRIME® transfection reagent (Polyplus) according to the manufacturer’s instruction. Lentiviral packaging plasmids pCMV-DR8.8 and pMD2.G were co-transfected with the backbone plasmid into 293T cells for virus production. Cells were selected in 2.5 µg/mL puromycin in the culture medium or by fluorescence-activated cell sorting to generate the stable transfections.

### Cell scratch wound healing assay

Cells were plated at a density of 1×10^5^ cells/well in triplicate into 6-well plates. Once the cells had spread over the bottom of the wells, three or four parallel lines were scratched into each well using sterile 10μL tips. Suspended cells were washed off and remaining cells were cultured in medium without FBS. After 12, 24 and 36 hours of incubation, for each well, five random fields were examined under a light microscope, photographed, and counted manually.

### Migration assay

Costar Trans-well migration plates with 8μm pore size (Corning, #3422) were pre-coated with Matrigel. Cells (1×105) in 100μL DMEM medium without FBS were placed in triplicate into the upper chamber. To the lower chamber, 500 μL medium containing 10% FBS was added. After 12, 24 and 36 hours of incubation, the plate inserts were removed and washed with PBS buffer several times to get rid of unattached cells. All the residual cells on the upper side were scraped with a cotton swab. Migrated cells on the lower side of the insert were fixed in 4% formalin for 20 minutes, washed twice with PBS, and stained with 0.1% crystal violet for 10 minutes. For each insert, five random fields were examined under a light microscope, photographed, and counted manually.

### Data mining using public database

The gene expression data for renal clear cell carcinoma (KIRC) was downloaded from TCGA, which were processed by Broad Institute’s TCGA workgroup. The RNA-seq level 3 gene expression data contain log2- or log10-transformed RNA-seq by expectation maximization (RSEM) values summarized at gene level.

### Statistics

Statistical evaluation was conducted using Student’s t-test. Multiple comparisons were analyzed first by one-way analysis of variance. The log-rank (Mantel-Cox) test was used for patient survival analysis. The Pearson correlation was used to analyze the strength of the association between expression levels of SETD2 and its related genes in patient samples. A significant difference was defined as P <0.05.

### Study approval

All mice were maintained in a specific-pathogen-free (SPF) facility and all experimental procedures were approved by the institutional biomedical research ethics committee at the Shanghai Institutes for Biological Sciences or Institute of Zoology, Chinese Academy of Sciences.

### Data availability

RNA-Seq and ChIP-Seq raw data have been deposited in the Gene Expression Omnibus (GEO) under accession number GEO: GSE125381 and GSE125528

## Author contributions

H.R. performed most of the experiments, analyzed the data and wrote the paper. X.L., M.L., J.L., W.F. and J.X. helped with the experiments. L.L. and W.Q. G. conceived the concept, designed the experiments and drafted and revised manuscript. All authors edited and approved the final manuscript.

## Acknowledgments

This study was supported by funds from Ministry of Science and Technology of the People’s Republic of China (2017YFA0102900 to W.Q.G.), National Natural Science Foundation of China (81772938 to L.L., 81872406 and 81630073 to W.Q.G.), State Key Laboratory of Oncogenes and Related Genes (KF01801 to L.L.), Science and Technology Commission of Shanghai Municipality (18140902700 and 19140905500 to L.L., 16JC1405700 to W.Q.G.), KC Wong foundation (to W.Q.G.) and Innovation Research Plan from Shanghai Municipal Education Commission (ZXGF082101 to L.L.). The study is also supported by Bio-ID Center, School of Biomedical Engineering, Shanghai Jiao Tong University.

