## Supplemental information for "SETD2 deficiency impairs β-catenin destruction complex to facilitate renal cell carcinoma formation"

### Supplemental data

Figure S1

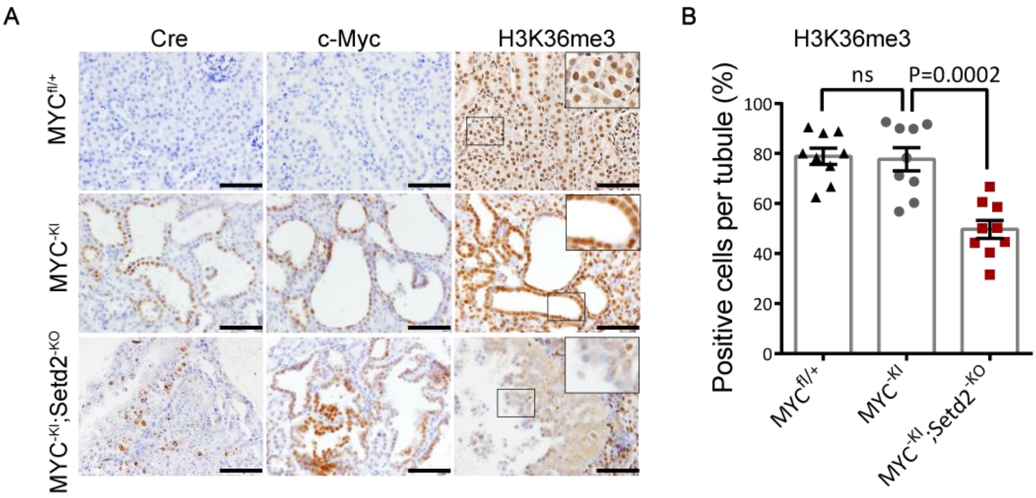


**Figure S1. c-MYC is overexpressed and Setd2 is knocked out in MYC$^{-KI}$ and MYC$^{-KI}$; SETD2$^{-KO}$ mice.** (A-B) Representative IHC images showing protein expression upon MYC activation and Setd2 deficiency. Scale bars: 80 μm in A. Statistical comparisons were made using a 2-tailed Student's t test. Data are represented as mean ± SEM.

### Figure S2

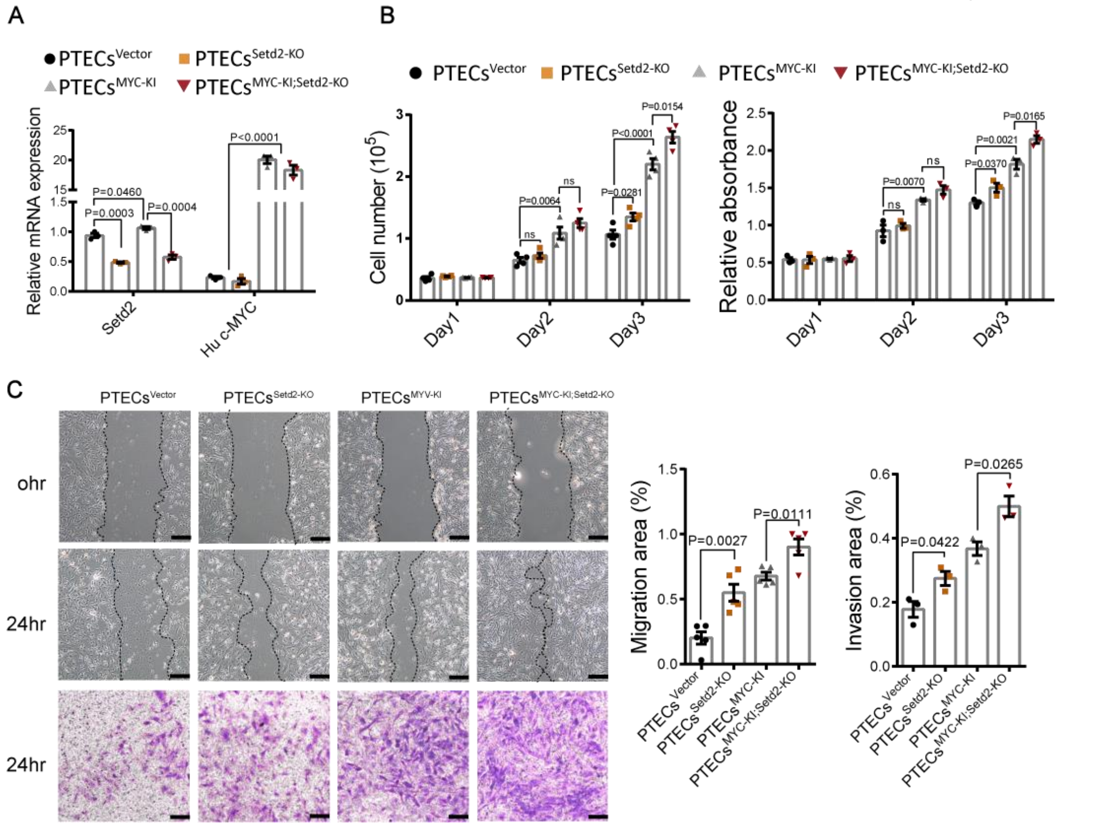


**Figure.S2 Setd2 deficiency promotes PTECs proliferation, migration and invasion *in vitro*.** (A) RT-qPCR analysis of $PTECs^{Vector}$, $PTECs^{SETD2-KO}$, $PTECs^{MYC-KI}$ and $PTECs^{MYC-KI;SETD2-KO}$ showing Setd2 and c-MYC expression level. (B) Cell proliferation abilities of $PTECs^{Vector}$, $PTECs^{SETD2-KO}$, $PTECs^{MYC-KI}$ and $PTECs^{MYC-KI;SETD2-KO}$. (C) Cell migration and invasion abilities of $PTECs^{Vector}$, $PTECs^{SETD2-KO}$, $PTECs^{MYC-KI}$ and $PTECs^{MYC-KI;SETD2-KO}$, as demonstrated by cell wound scratch and trans-well assays. Scale bars: 80 μm in C. Statistical comparisons were made using a 2-tailed Student's t test. Data are represented as mean ± SEM.

### Figure S3

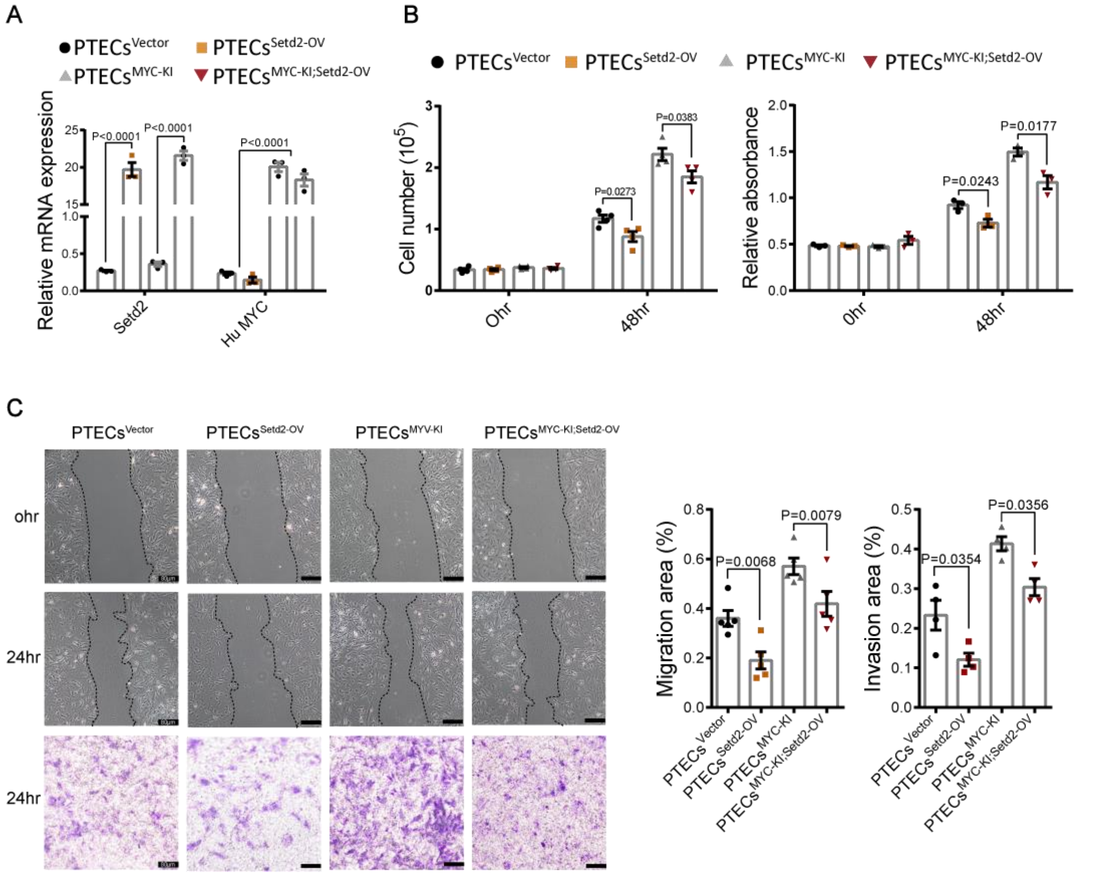


**Figure.S3 Setd2 overexpression inhibits PTECs proliferation, migration and invasion *in vitro*.** (A) RT-qPCR analysis of PTECs$^{Vector}$, PTECs$^{SETD2-OV}$, PTECs$^{MYC-KI}$ and PTECs$^{MYC-KI;SETD2-OV}$ showing Setd2 and c-MYC expression level. (B) Cell proliferation abilities of PTECs$^{Vector}$, PTECs$^{SETD2-OV}$, PTECs$^{MYC-KI}$ and PTECs$^{MYC-KI;SETD2-OV}$. (C) Cell migration and invasion abilities of PTECs$^{Vector}$, PTECs$^{SETD2-OV}$, PTECs$^{MYC-KI}$ and PTECs$^{MYC-KI;SETD2-OV}$, as demonstrated by cell wound scratch and trans-well assays. Scale bars: 80 μm in C. Statistical comparisons were made using a 2-tailed Student's t test. Data are represented as mean ± SEM.

Figure S4

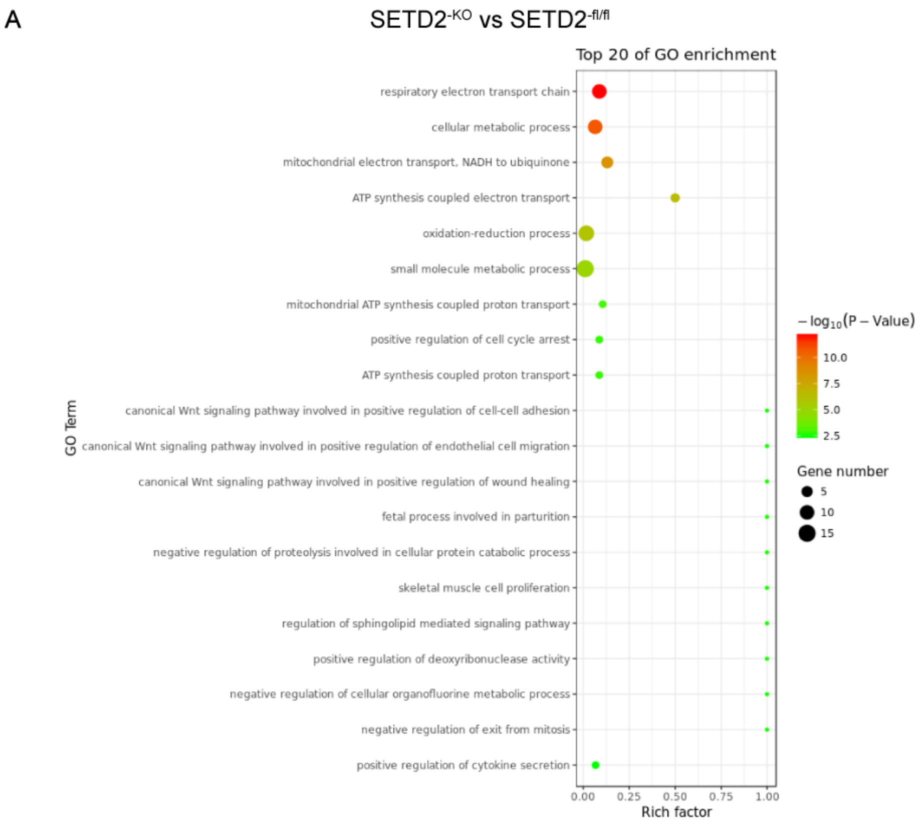


**Figure.S4 Setd2 deficiency enriched the gene concepts linked to hyperactive Wnt/β-catenin signaling.** (A) The genes that were significantly, differentially expressed in Setd2 deficient kidneys were tested for enrichment and represented using ClueGO.

Figure S5

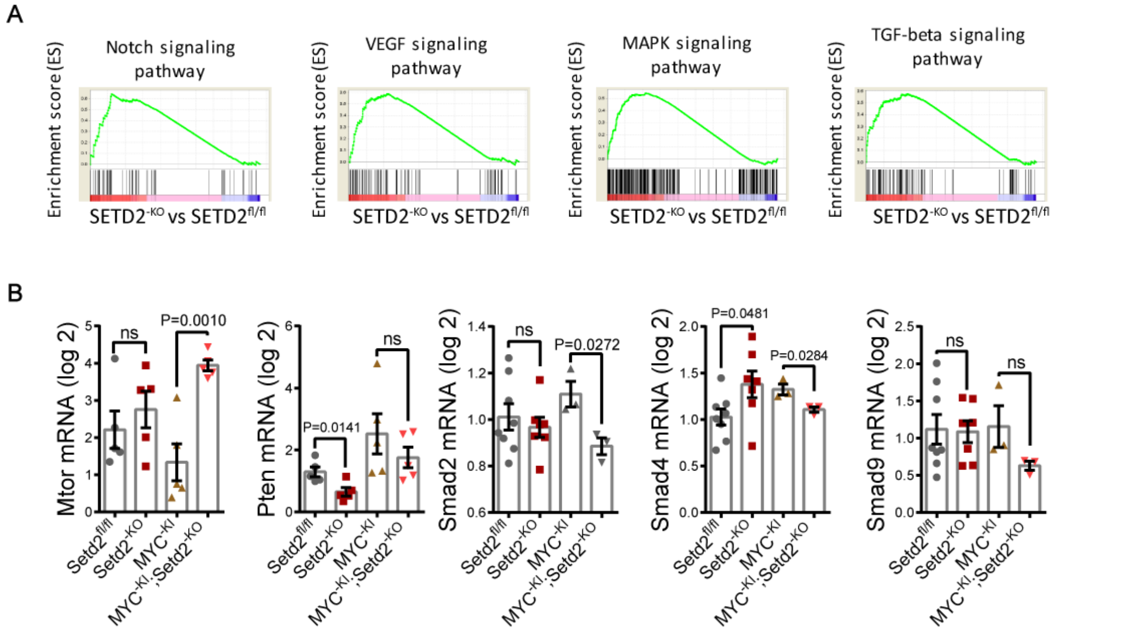


**Figure.S5 GSEA enrichment plots of different pathways.** (A) GSEA enrichment plots of differentially expressed genes belonging to Notch signaling, VEGF signaling, MAPK signaling and TGF-β signaling associated with SETD2 deletion. (B) Expression levels of some markers of different signaling pathways associated with SETD2 deletion. Statistical comparisons were made using a 2-tailed Student's t test. Data are represented as mean ± SEM.
